# The pericardium promotes cardiac repair and remodelling post-myocardial infarction

**DOI:** 10.1101/771154

**Authors:** Katie J. Mylonas, Lucy H. Jackson-Jones, Jack P. M. Andrews, Marlene S. Magalhaes, Marco Meloni, Nikhil V. Joshi, Judith E. Allen, David E. Newby, Marc R. Dweck, Gillian A. Gray, Cécile Bénézech

## Abstract

The pericardium is widely recognised for its lubricating and bio-mechanical properties. It also contains fat-associated lymphoid clusters (FALCs) and its immune functions have been widely overlooked. Here we aimed to assess the inflammatory activity of the pericardium in patients who suffered a recent myocardial infarction (MI) and to determine its importance for repair and remodelling in a murine MI model induced by coronary artery ligation (CAL). By comparing ^18^F-fluorodeoxyglucose (FDG) activity in the pericardium of patients with stable coronary artery disease and patients who had a recent MI, we demonstrate that MI is associated with increased pericardial inflammation. We confirm in mice, that pericardial FALCs undergo a major expansion following CAL. We show that despite similar initial injury, removal of the pericardium prior to MI disrupted subsequent repair, resulting in 50% mortality due to cardiac rupture, while all mice with intact pericardia survived. Removal of the pericardium also led to decreased staining for Ym1, a marker of reparative macrophages and adverse cardiac fibrosis within the infarct area. Together, this work indicates a crucial role for the pericardium in regulating inflammation, macrophage polarisation and tissue remodelling in the heart following MI.

## Introduction

Complications of acute myocardial infarction (MI), including rupture and the development of heart failure, remain a major cause of death and disability. The inflammatory response initiated after MI is intricately linked to the extent of the injury, the quality of repair and the level of remodelling of the myocardium.^1^ While the recruitment of neutrophils is clearly associated with a robust inflammatory response and contribute to heart failure after ischemic heart injury,^2^ the role of macrophages remains more ambiguous. Macrophages are heterogenous in origin and function, and contribute to both the early inflammatory response and the repair and remodelling phase.^3, 4^ Better understanding of the mechanisms controlling inflammation and macrophage recruitment following MI is thus vital to the development of new and effective treatments.^1^

The pericardium is a double-layered fluid-filled serous cavity that protects and anchors the heart. It is delimited by two membranes, the visceral pericardium (or epicardium) lining the surface of the heart and the parietal pericardium lining the outer layer of the pericardial cavity. In man, both membranes incorporate adipose tissue covered with a mesothelial monolayer. In mouse, only the parietal pericardium contains adipose tissue, the epicardium being only made of a monolayer of mesothelial cells. We recently showed that the adipose tissue of the pericardium contains numerous small immune structures called fat-associated lymphoid clusters (FALCs).^5, 6^ These are critical for the maintenance and function of the B cells of the serous cavities^7-9^ which represent a unique compartment of B cells specialised in the production of natural antibodies important for homeostasis and protection against infection.^10^ During peritonitis, the number and size of FALCs of the omentum and mesenteries increases and FALCs recruits inflammatory cells.^5, 9^ Recent work in mice by Horckmans *et al*. demonstrated a strong correlation between pericardial FALC B cell activation, the extent of cardiac inflammation and functional outcomes following MI.^11^ Using cannabinoid receptor CB2 deficient mice (CB2^-/-^) which have increased levels of circulating B cells, they concluded that secretion of Granulocyte-macrophage colony-stimulating factor (GM-CSF) by pericardial B cells promotes bone marrow granulopoiesis, cardiac neutrophil infiltration, enhances fibrosis and worsens functional cardiac outcome.^11^ Furthermore, pericardial cavity macrophages have a functional phenotype comparable to other F4/80^hi^ serous cavity macrophage populations and are controlled by the expression of the transcription factor GATA-6.^12-14^ Deniset *et al* have shown that these macrophages relocate to the injured heart where they act to suppress cardiac fibrosis.^15^ This led us to ask whether the pericardium was also involved in the regulation of the inflammatory response after MI in man and in mice with a normal B cell compartment.

Here we demonstrate that MI is associated with increased pericardial inflammation in patients who have suffered a recent MI. We provide direct evidence in mice, that MI leads to a pericardial inflammatory response which mirrors that of the infarct. Furthermore, we show that the pericardium promotes healing and survival after MI and suggest this occurs via the redistribution of serous cavity macrophages to pericardial FALCs which contribute to the remodelling of the collagen matrix within the infarct zone.

## Materials and Methods

### Clinical assessment of pericardial ^18^F-FDG uptake

This study was done with the approval of the local research ethics committee, in accordance with the Declaration of Helsinki, and with informed consent of each participant.^16^ ^18^F-FDG cardiac PET-CT scans from 37 patients with stable coronary artery disease (angina) and 36 patients with unstable coronary artery disease (recent myocardial infarction), originally recruited for a ^18^F-FDG and ^18^F-NaF comparison study described by *Joshi et al*., were retrospectively analysed.^16^ Patients were predominantly middle-aged men with multiple cardiovascular risk factors. To minimise myocardial uptake, all patients were instructed to adhere to a low carbohydrate, high protein and high fat diet for at least 24 hours prior to their ^18^F-FDG PET-CT scan. PET-CT’s were performed on average 8 (IQR 3-10) days after symptom onset in the unstable group. Electrocardiograph-gated PET images were reconstructed in diastole (50–75% of the R-R interval, Ultra-HD) using the Siemens Ultra-HD algorithm, fused with the non-contrast CT, and analysed by an experienced observer (JA). Images were analysed using an OsiriX workstation (OsiriX version 5·5·1 64-bit; OsiriX Imaging Software, Geneva, Switzerland). Patients underwent ^18^F-FDG (90 min after 192 MBq) PET-CT scanning within a median of 6 (IQR 3–9) days. Predefined myocardial suppression of ^18^F-FDG uptake was achieved in 70% of patients (median myocardial standard uptake value 3·92 [IQR 2·71–5·55]). Two-dimensional 50 mm^2^ circular regions of interest were drawn within the epicardial tissue within the right coronary sulcus, maximising the distance from the right coronary artery and reducing potential problems with spill over of signal. The maximum standard uptake value (the decay corrected tissue concentration of the tracer divided by the injected dose per bodyweight) was measured and corrected for mean blood pool activity in the right atrium to provide the maximum tissue-to-background ratio (TBRs) measurements. It was not possible to standardise the size and shape of ROI drawn within these anatomical structures and so mean SUV and TBR values were not measured.

### Mice and CAL surgery

All animal work was compliant with IACUC guidelines, conducted in accordance with the UK Government Animals (Scientific Procedures) Act 1986 and was approved by the University of Edinburgh Animal Welfare and Ethical Review Board. Aged-matched C57BL/6 male mice (6-10 weeks old) were used for all experiments. CAL surgery was carried out as previously described.^17^ The fibrous-pericardium was either fully removed or left ‘intact’ during this procedure. In ‘intact’ mice, a small snip was made in the fibrous-pericardium to expose the coronary artery for ligation, but it was otherwise kept in one piece. Alternatively, the fibrous-pericardium was removed, placed in RPMI media while the CAL surgery took place and then engrafted back onto the heart, where it adhered. For plasma troponin measurement, tail vein blood was collected from mice 24 hours post-CAL surgery. Cardiac Troponin-I (Tn-I) was measured in the plasma using the mouse high sensitivity Tn-I ELISA kit according to the manufacturer instructions (Life Diagnostics, UK). Any deaths were recorded daily. Post-mortem examination confirmed deaths occurred following heart-rupture. Mice were infected sub-cutaneously with 25–30 *Ls* L3’s.

### Optical projection tomography (OPT)

6 hours after surgery, hearts were recovered and processed for imaging and scanned in a calibrated Bioptonics 3001 tomograph.^18^ In the Cy3 fluorescence emission channel (545nm) the viable myocardium had strong autofluoresence (dark) and the injured myocardium appeared lighter. The suture placement of each was also clearly visible. Tomographic reconstruction using Nrecon software produced hundreds of image files depicting vertical transverse slices through the heart. These were analysed using Analyze12 software allowing delineation between viable (dark) and injured myocardium (light areas). This allowed quantification of total % injury between intact and injured left ventricles. Noting the slice wherein the suture became apparent also allowed for analysis of suture placement between both groups.

### Immunohistochemistry (IHC)

Formalin-fixed paraffin-embedded heart sections were blocked then incubated overnight at 4°C with primary antibody. Secondary antibody was then added for 1 hour and revealed using Peroxidase-labelled ABC reagent DAB substrate (Vector Laboratories, UK). See antibody list. Finally, the sections were counterstained with haematoxylin, dehydrated through ethanol and xylene and mounted with DPX mountant (Sigma). For picro-sirius red (PSR) staining of collagen, sections were deparaffinised and rehydrated before treatment in haematoxylin for 8 minutes and stained in PSR for 1 hour. For quantification, sections were tiled at ×40 magnification (Image Pro6.2, Stereologer Analyser 6 MediaCybernetics). The % area stained was calculated within the infarct and border area.

Wholemount immunofluorescence (IF) staining and confocal microscopy Pericardium samples were fixed for one hour on ice in 10% NBF (Sigma) and then permeabilized in PBS 1% Triton-X 100 (Sigma) for 20 minutes at room temperature prior to staining with primary antibodies for one hour at room temperature in PBS 0.5% BSA 0.5% Triton. After washing in PBS, tissues were stained with secondary antibodies for one hour at room temperature in PBS 0.5% BSA 0.5% Triton. For neutral lipid staining, fixed pericardia were incubated with LipidTox (Invitrogen) for 20 minutes at room temperature prior to staining with primary antibodies. Antibodies used are listed in Supplemental Table 1. After mounting with Fluoromount G, confocal images were acquired using a Leica SP5 laser scanning confocal microscope. Image 3D reconstruction was created using Fiji.

### Flow cytometry

The pericardia were enzymatically digested with 1mg/ml Collagenase D (Roche) for 35 minutes at 37°C in RPMI 1640 (Sigma) containing 1% Fetal Bovine Serum (Sigma). Heart tissue (infarct and surrounding border) was digested in Collagenase II and DNase 1 (600 U/ml Collagenase II; CLS-2 Worthington, 60 U/ml DNAse 1; Ambion, Warrington) in Hank’s Balanced Salt Solution (HBSS; GIBCO) at 37°C for 30 min following dissociation by GentleMacs dissociator (Miltenyl). Cells were stained with LIVE/DEAD (Invitrogen), blocked with mouse serum and anti-murine CD16/32 (clone 2.4G2, Biolegend) and stained for cell surface markers (See Supplemental Table 1 for list of antibodies used). All samples were acquired using a BD Fortessa and analyzed with FlowJo software (Tree Star).

### Statistics

All values are expressed as mean ± SEM. Unpaired Student’s t-test or ANOVA with post-hoc test were used for analysis. A X^2^ (Chi-square) test was used for Kaplan-Meier survival curve analysis. P-values <0.05 denote statistical significance, *p<0.05, **p<0.01, ***p<0.005.

## Results

Pro-inflammatory macrophages and rapidly proliferating cells have a high metabolic rate and avidly accumulate glucose.^19, 20^ To determine if MI had an impact on the inflammatory activity of the pericardium, we retrospectively analysed ^18^F-FDG cardiac PET-CT scans from patients with stable coronary artery disease vs patients with recent MI.^16^ Two-dimensional 50 mm^2^ circular regions of interest within the epicardial tissue of the right coronary sulcus were drawn, maximising the distance from the right coronary artery (**Figure 1A)**. The maximum standard uptake value (SUVmax) was measured and corrected for mean blood pool activity in the right atrium to provide the maximum tissue-to-background ratio (TBRmax) measurements. Pericardial ^18^F-FDG uptake increased in patients following myocardial infarction compared to patients with stable coronary artery disease (SUVmax 1.42 vs 1.11, p = 0.0053; TBRmax 1.35 vs 0.80, p=0.0001) (**Figure 1B)**. This was despite patients in the stable cohort having a higher burden of atherosclerotic plaque than patients in the MI cohort (599 [IQR 60–1302] *vs* 159 [42–456] Agatston units, p=0·006).

**Figure 1:**
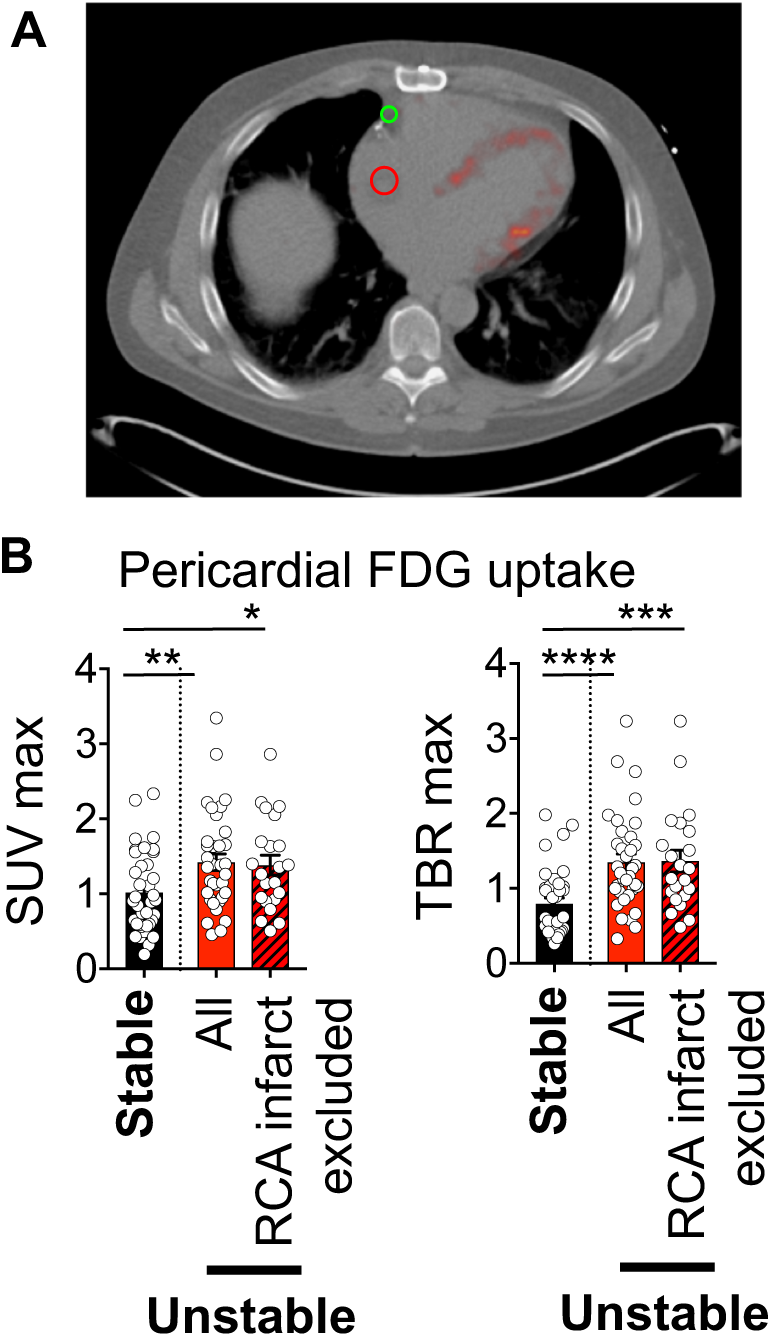
The uptake of ^18^F-FDG by the pericardium is increased in patients who had a recent MI. **A:** Representative electrocardiograph-gated PET image showing the circular regions drawn within the epicardial tissue within the right coronary sulcus (green circle) to determine pericardial SUVmax and within the right atrium (red circle) to determine the blood pool activity and pericardial TBRmax. **B:** Pericardial SUVmax and TBRmax calculated in patients with stable coronary heart diseases (n=37) and unstable patients (with all unstable patients included n=35, red bar; and with patients with RCA territory infarcts excluded n=22, red bar with black hashing). Mann Whitney test were applied, **=P <0.01, *** P=<0.001, **** P=<0.0001.

Uptake of ^18^F-FDG has previously been shown to be elevated in ruptured and inflamed atherosclerotic plaque.^20^ As the regions of interest were sampled from within the right coronary sulcus, culprit plaques within the right coronary artery could potentially influence the accuracy of pericardial sampling. To improve the validity of our results, a further analysis was therefore performed to exclude those patients with right coronary artery (RCA) territory infarcts. In patients having had an MI involving the left coronary artery (n = 14), pericardial 18F-FDG uptake was again increased compared to the patients with stable coronary artery disease (SUVmax 1.38 vs 1.02, p = 0.0197; TBRmax (1.36 vs 0.80, p=0.0003) **(Figure 1B).**

To prove that MI induced an inflammatory response in the pericardium, we investigated the effect of CAL-induced MI on the immune composition of the parietal pericardium and FALCs in mice. MI led to an enlargement of the parietal pericardium which adhered spontaneously to the infarct zone (**Figure 2A**). The weight of the pericardium increased 2-fold 3 days after MI, confirming that MI led to expansion of the pericardium (**Figure 2B**). MI led to a massive enlargement of FALCs as determined by staining for the hematopoietic marker CD45 and the B cell marker IgM, while the adipose tissue stained with the neutral lipid marker Lipidtox, decreased (**Figure 2C**). CAL-induced MI led to a rapid recruitment of neutrophils and monocytes into the infarct area that was detected 24 hours after MI and was followed by the recruitment of F4/80^+^ macrophages from day 2 after MI (**Figure 2E** and **Supplementary Figure 1**). We detected a concomitant recruitment of neutrophils and monocytes in the parietal pericardium. The same number of neutrophils and monocytes were recruited into the infarct area and the parietal pericardium indicating that the recruitment of inflammatory cells observed in the pericardium was quantitively not negligible when compared to the infarct area (**Figure 2D, 2E** and **Supplementary Figure 1**). In addition, MI led to a rapid recruitment of F4/80^+^MHCII^-^ macrophages into the parietal pericardium, which was detectable as early as 24 hours following MI suggesting that the recruitment of macrophages to the pericardium preceded the recruitment of macrophages into the infarct area (**Figure 2D, 2E**). Macrophages recruited in the parietal pericardium lacked expression of MHCII despite high expression of F4/80, in contrast to the MHCII^+^ macrophages found in the parietal pericardium of naïve mice Figure 2D). Lack of MHCII expression is typical of serous cavity macrophages^21, 22^ and MHCII expression did not increase on pleural cavity macrophages across the CAL time course (**Supplementary Figure 2**). This suggested that recruited pericardial macrophages came from the pericardial or pleural cavities. The recruitment of CD11b^+^F4/80^high^ macrophages in the parietal pericardium was concentrated in FALCs 3 days after MI (**Figure 2F**). In our previous work, we observed that infection with the filarial nematode *Litomosoides sigmodontis* (Ls), which resides in the pleural cavity, led to activation of pericardial FALC B cells,^6^ but did not induce the recruitment of resident pleural F4/80^high^ macrophages into pericardial FALCs (**Supplemental Figure 3**). Recruitment of serous macrophages into pericardial FALCs thus appears to be specific to heart injury. Taken together, these data indicated that MI led to an important remodelling of the pericardial adipose tissue with FALCs increasing dramatically in size, the entrance of neutrophils and monocytes as well as the recruitment of serous macrophages into pericardial FALCs.

**Figure 2:**
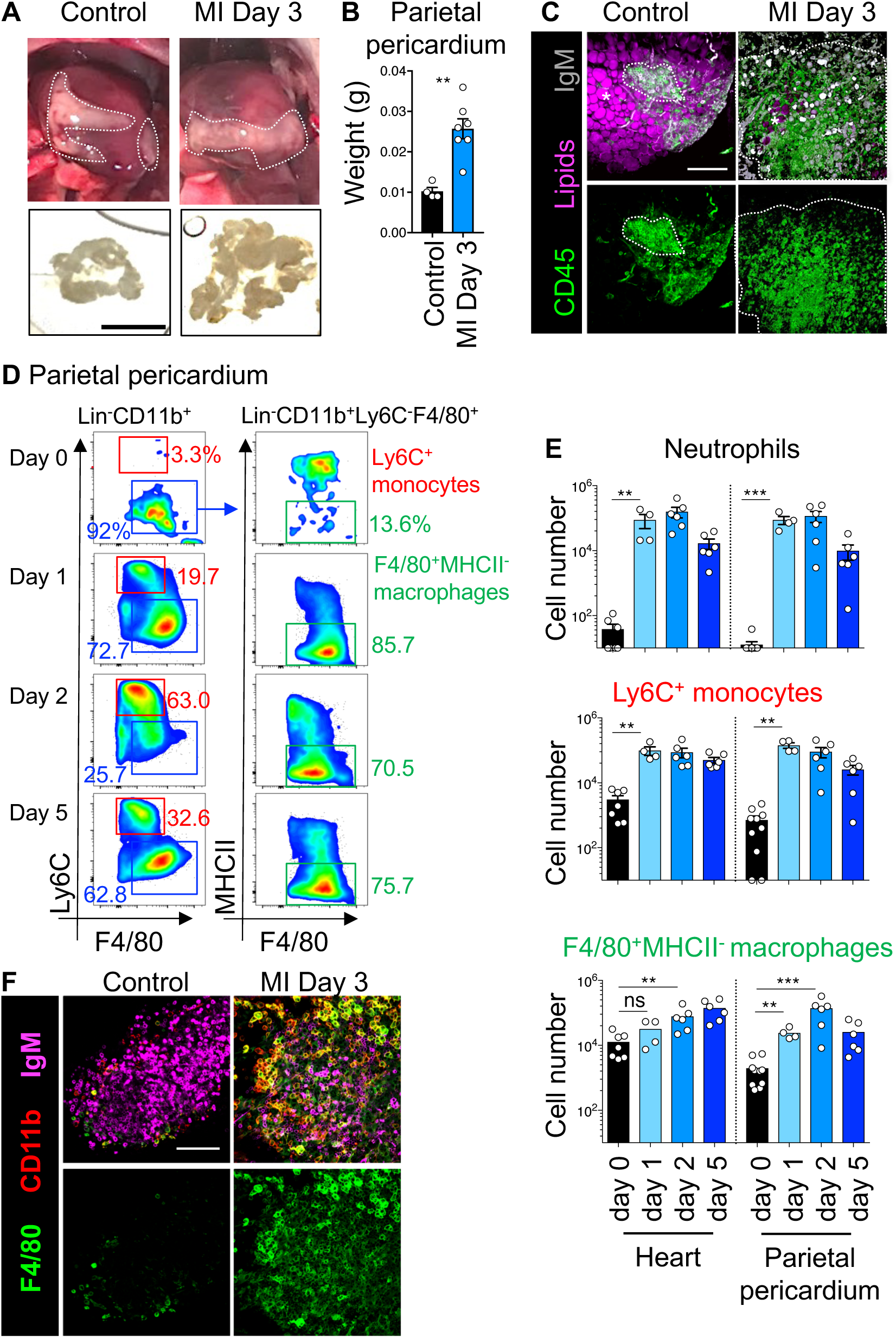
CAL-induced MI triggers an inflammatory response in the pericardium. **A:** Macroscopic picture of the pericardium and heart in a control mouse and a mouse 3 days post-CAL (upper panels). Lines highlight the adipose tissue of the parietal pericardium. Isolated parietal pericardium from the same mouse (lower panels, scale bar 1 cm). **B:** Weight of the pericardium isolated. Data pooled from two independent experiments n=4-7 mice per group. **C:** Representative 3D reconstruction of FALCs obtained by confocal imaging of wholemount immunofluorescence staining of pericardia from control mice and mice 3 days post-MI. Hematopoietic cells stained with CD45 (green), lipids stained with LipidTox (purple) and B cells stained with IgM (grey). Scale bar 100 µm. **D:** Representative flow-cytometric analysis used to identified monocytes (red gate) and macrophages (blue gate) in the parietal pericardium of a naïve mouse (Day 0) and mice at Day 1, 2 and 5 post CAL. **E:** Number of neutrophils, monocytes and macrophages found in the infarct area and pericardium of control mice and mice at day 1, 2 and 5 post-MI. Data pooled from at least two independent experiments n=4-10 mice per group. See additional gating strategy in Supplemental Figure 1. **F**: Representative confocal images of wholemount immunofluorescence staining of pericardia from control mice and mice 3 days post-MI. Myeloid cells stained with CD11b (red), macrophages stained with F4/80 (green) and B cells stained with IgM (purple). Scale bar 50 µm. Mann Whitney test were applied, ns= non-significant, **=P <0.01, *** P=<0.001.

Since the pericardium was involved in the inflammatory response induced by MI, we assessed the importance of this tissue for recovery and healing following MI. We performed CAL or sham surgery with the membranes of the parietal pericardium either completely removed or minimally disrupted during surgery. The rupture rate in male C57BL/6 mice after conventional MI surgery with disrupted pericardium is typically of 25%.^23-26^ Keeping the pericardium intact during surgery led to 100% survival. In contrast, complete removal of the pericardium resulted in death of 50% of mice by rupture, mostly between days 2-7 after MI. Mice with ‘intact’ pericardia, and sham operated mice all survived (**Figure 3A**). This was despite similar initial injury in all groups, as measured by plasma troponin concentrations at 24 hours (**Figure 3B**) and by calculating the % left ventricular injury by optical projection tomography (OPT) performed 6 hours post-MI (**Figure 3C**).^18^ OPT analysis also confirmed that the suture placement during CAL surgery was uniform whether the fibrous-pericardium was present or not (**Figure 3C**). To rule out the possibility that mechanical damage caused by removal of the pericardium was the cause of the rupture, CAL surgery was undertaken with membranes of autologous pericardium grafted back onto the heart. Examination of the heart 7 days after MI showed that the grafted pericardial membranes remained in place, covering the infarcted area (**Figure 3D**). Grafting the parietal pericardium back on the heart was sufficient to restore survival (**Figure 3E**), indicating that the parietal pericardium was protective post-MI, and that disruption of the pericardial cavity as such was not the cause of heart rupture post-CAL.

**Figure 3:**
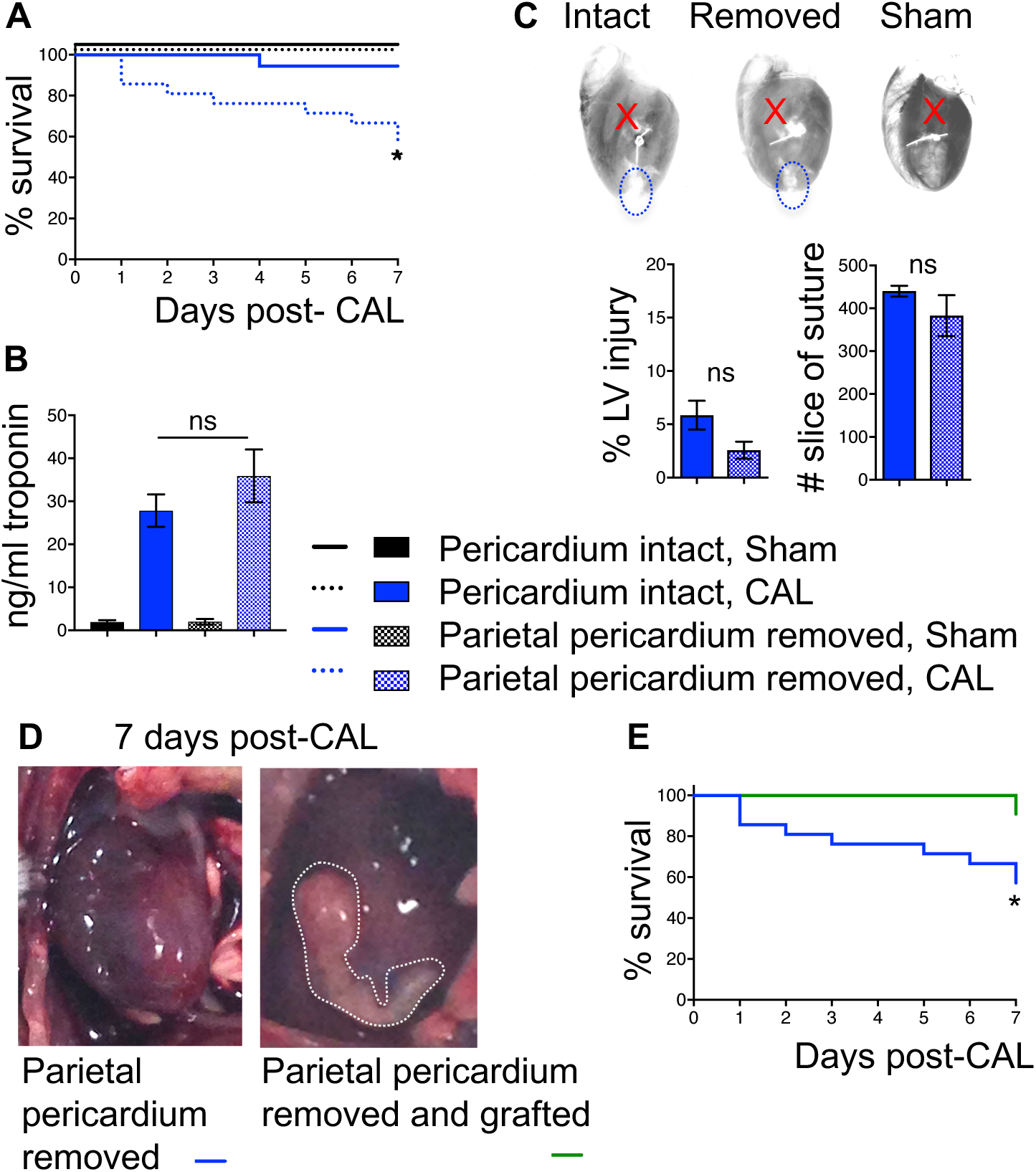
Removal of the pericardium results in increased death post-MI despite similar initial injury. **A-C**: The pericardium was either removed or left intact prior to CAL or sham (no ligation of coronary artery) surgeries. Survival curve at 7 days (**A**). Plasma troponin levels 24 hours post-MI (**B**) Representative heart OPT images showing infarct area (dotted circles) and suture placement (red cross) at 6 hours post-CAL (upper row). Quantification of the total % injury between intact and injured left ventricle (LF). Quantification of the placement of the suture as assessed by the optical slice wherein the suture became apparent (**C**). **D-E:**The pericardium was removed at time of surgery and then re-grafted back onto the heart. Macroscopic picture of the pericardium and heart in a mouse with the pericardium removed and a mouse with the pericardium removed and re-grafted at 7 days post-MI (**D**). Survival curve at 7 days. Note that data in A and E used the same cohort of mice which underwent CAL with pericardium removed. Data pooled from at least two independent experiments: n=8 (Pericardium intact, Sham), n=18 (Pericardium intact, CAL), n= 16 (Parietal pericardium removed, Sham), n=21 (Parietal pericardium removed, CAL), n=11 (Parietal pericardium removed and grafted CAL) mice per group. For Kaplan-Meier survival curve analysis, a X^2^ test was used, *=P <0.05. For troponin assays Kruskal Wallis test with Dunn’s multiple comparisons test was performed. For LV injury and slice of suture measurements, Mann Whitney tests were applied. ns= non-significant.

We next investigated whether removal of the parietal pericardium led to a change in the recruitment of macrophages in the infarct area. Ym1 (Chil3) is highly expressed by M2-like reparative macrophages in post-MI heart.^24^ We found that removal of the fibrous pericardium was associated with decreased IHC staining for Ym1 in the heart 7 days after MI (**Figure 4A**). Furthermore, grafting of the pericardium was sufficient to restore Ym1 staining to levels comparable to mice with an intact pericardium. The decrease in Ym1 staining was only apparent in the sub-epicardial layer, where the parietal pericardium was adhering to the infarct zone suggesting that the close proximity of the pericardium directly influenced the polarisation of macrophages toward a reparative phenotype or that Ym1^+^ reparative macrophages migrate from the pericardium to the infarct (**Figure 4A**). M2-like reparative macrophages are required for the early deposition of collagen.^24^ We thus assessed whether removal of the pericardium would be associated with decreased deposition of collagen after MI as measured by Picro Sirius Red (PSR) staining. However, removal of the pericardium led to increase levels of collagen in the infarct area at day 7 after MI compared to mice with an intact pericardium or mice with a grafted parietal pericardium, (**Figure 4 C-D**). There was a similar pattern in the remote myocardium showing increased collagen deposition throughout the rest of the tissue in absence of the pericardium (**Supplemental Figure 4**). Our result thus suggests that the pericardium limits adverse fibrosis in the infarct and surrounding healthy tissue at early time point after MI.

**Figure 4:**
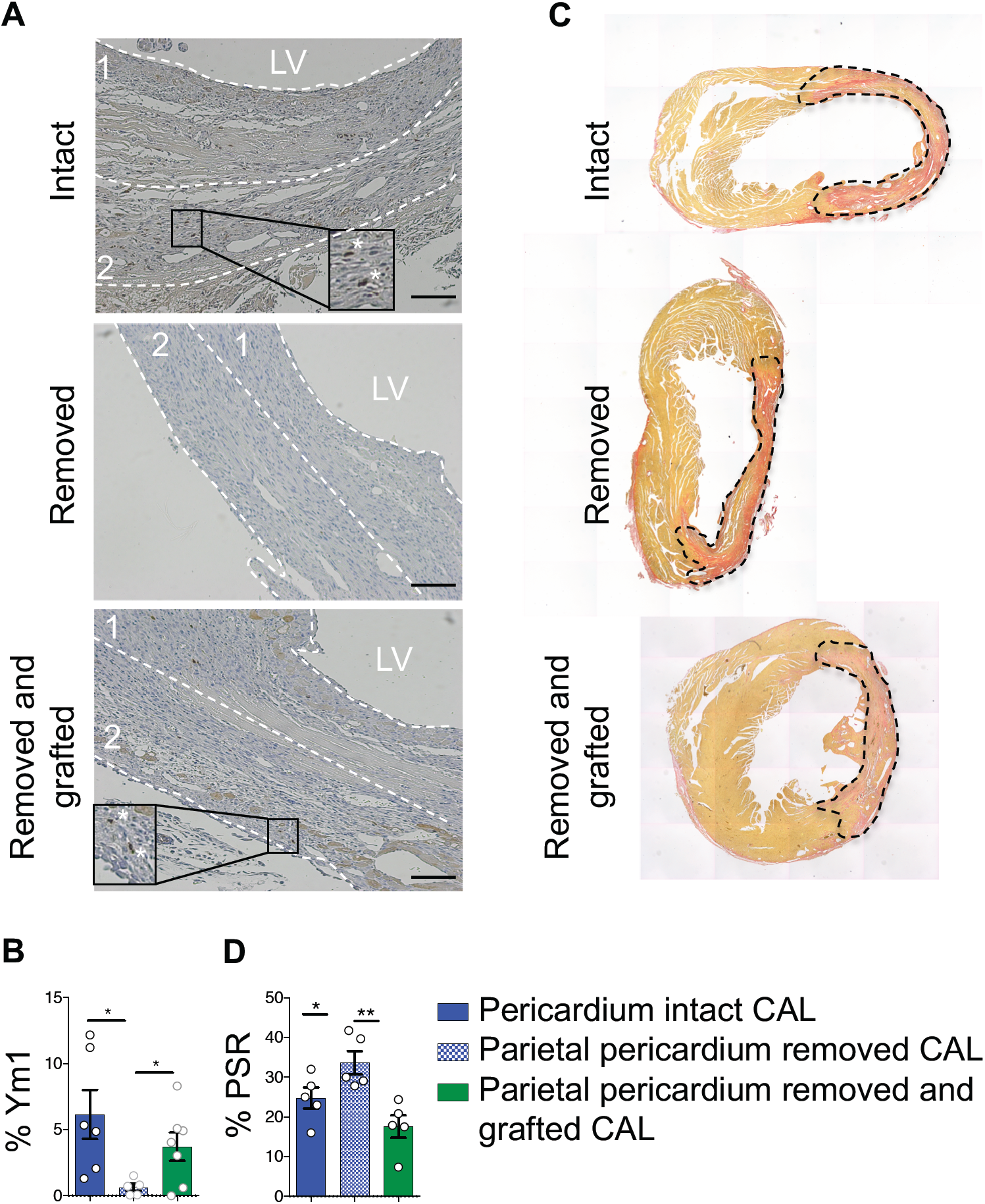
Removal of the pericardium results in decreased Ym1 staining and increased PSR staining in the infarct zone. The pericardium was either left intact, removed or removed and re-engrafted during CAL surgeries. Section of hearts were analyzed at day 7 post-MI. **A:** Representative micrographs within the left ventricular free wall showing Ym1 staining (brown). Dashed lines represent divisions between the subepicardial and subendcocardial regions. **B**: Quantification of % positive Ym1 staining in the subepicardial region. Data pooled from two independent experiments n=5-7 mice per group. Scale bar: 100 µM. **C**: Representative micrographs within the left ventricular free wall showing PSR staining (red) and **D**: quantification of % positive PSR staining. Data pooled from two independent experiments n=5 mice per group. Unpaired student’s T-test were performed. *=P <0.05, ** P=<0.01.

## Discussion

Here we demonstrate for the first time in human that MI is associated with increase inflammatory activity of the pericardium and we show in a murine model of MI that pericardial FALCs are involved in the inflammatory response following MI and that the parietal pericardium is critical for survival and the early recruitment of reparative macrophages and remodelling of the infarct tissue post-MI (**Figure 5**). Combined with recent published work,^11,15^ our data suggest that the pericardium represents a physiological route for cell recruitment into the heart and that FALCs play a critical role in the regulation of the inflammatory response within the heart.

**Figure 5:**
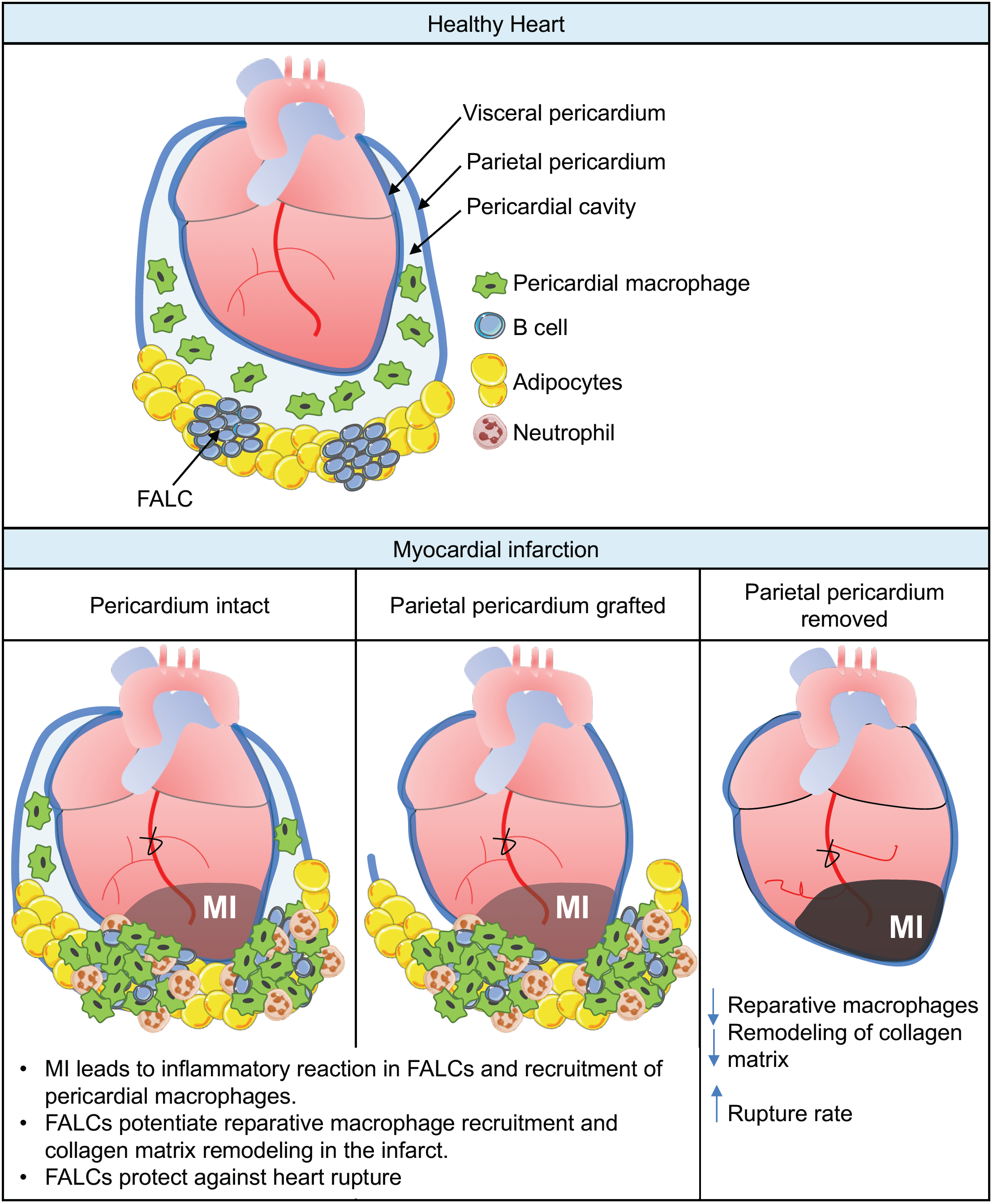
Proposed model.

While rupture is relatively rare complication of MI in patients, in the mouse it is typically a sign of disrupted repair and adverse inflammation^24, 27^ which can be caused by dysregulated recruitment of inflammatory cells into the heart post-MI. Horckmans et al. reported that the adverse pericardial inflammation driven by pericardial B cells in CB2^-/-^ mice was deleterious after MI and that removal of the pericardium restored repair and heart function.^11^ However, they did not report increased rupture after MI in their WT mice, when removing the pericardium. The main difference between their experiments and ours, is their use of female mice which are less prone to rupture than male mice, which could have hidden the phenotype we observed.

Shiraishi *et al.* showed that intravenous injection was not an efficient route for heart engraftment of M2-polarized BMMϕ, but intra-pericardial injection in a matrigel solution allowed their successful engraftment in the heart, improving survival and repair after MI in mice deficient in reparative Mϕ, further highlighting the physiological importance of the pericardium as a cellular gateway to the heart.^24^ Changes in the immune status of the pericardium, such as induced by airway infections^6, 7^ are likely to affect the inflammatory response in the infarct following MI. Since there is an elevated incidence of MI during the first week following acute airway infection,^28-32^ understanding how airway infections impact on the protective function of the pericardium after MI will be of particular clinical importance.

## Acknowledgements

We are extremely grateful to the CALM facility for advice on flow cytometry and confocal microscopy.

## Notes

Source of support: Medical Research Council (MRC) UK Grant to CB (MR/M011542/1) and JEA (MR/K01207X/1), Tenovus Scotland small project grant to CB and KJM and Wellcome Trust ISSF2 to CB and KJM.

The authors have declared that no conflict of interest exists.

