## Supplemental Figures and Tables for "The pericardium promotes cardiac repair and remodelling post-myocardial infarction"

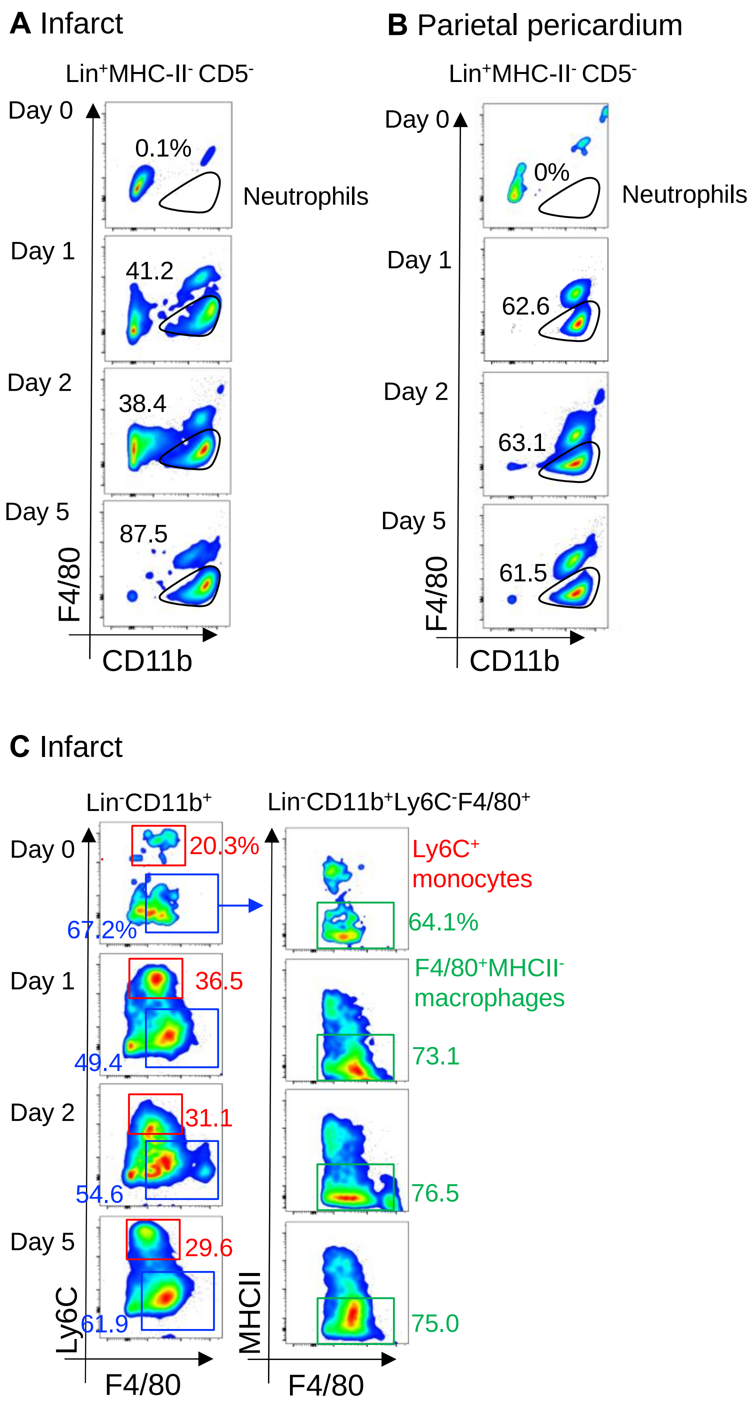


***Supplemental Figure 1: Gating strategy used to identified neutrophils, monocytes and macrophages in the infarct area and the parietal pericardium.*** **A,B:** Representative flow-cytometric analysis used to identified neutrophils in the infarct area (A) and parietal pericardium (B) of a naïve mouse (Day 0) and mice at Day 1, 2 and 5 post CAL. The lineage contained TCRβ, CD19, SiglecF and Ly6G. Lin+ cells were gated on and T cells and B cells were excluded using CD5 and MHCII markers. Eosinophils expressing low levels of F4/80, neutrophils were than identified as Lin+CD5-MHCII-F4/80- cells (black gate). **C**: Representative flow-cytometric analysis used to identified monocytes and macrophages in the infract area. Monocytes were identified as Lin-CD11b+Ly6C+F4/80- cells (red gate). Macrophages were identified as Lin-CD11b+Ly6C-F4/80+ cells (blue gate) and F4/80+MHCII+ macrophages were identified according to MHCII expression (green gate, second row).

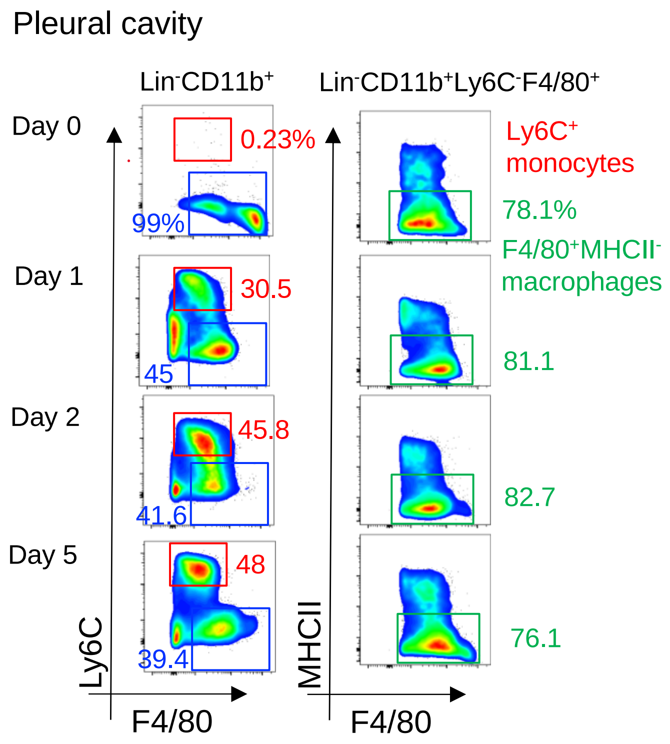


***Supplemental Figure 2: Gating strategy used to identified monocytes and macrophages in the pleural cavity.*** Representative flow-cytometric analysis used to identified monocytes and macrophages in the pleural cavity of a naïve mouse (Day 0) and mice at Day 1, 2 and 5 post CAL. Monocytes were identified as $Lin^{-}CD11b^{+}Ly6C^{+}F4/80^{-}$ cells (red gate). Macrophages were identified as $Lin^{-}CD11b^{+}Ly6C^{-}F4/80^{+}$ cells (blue gate) and $F4/80^{+}MHCII^{+}$ macrophages were identified according to MHCII expression (green gate, second row).

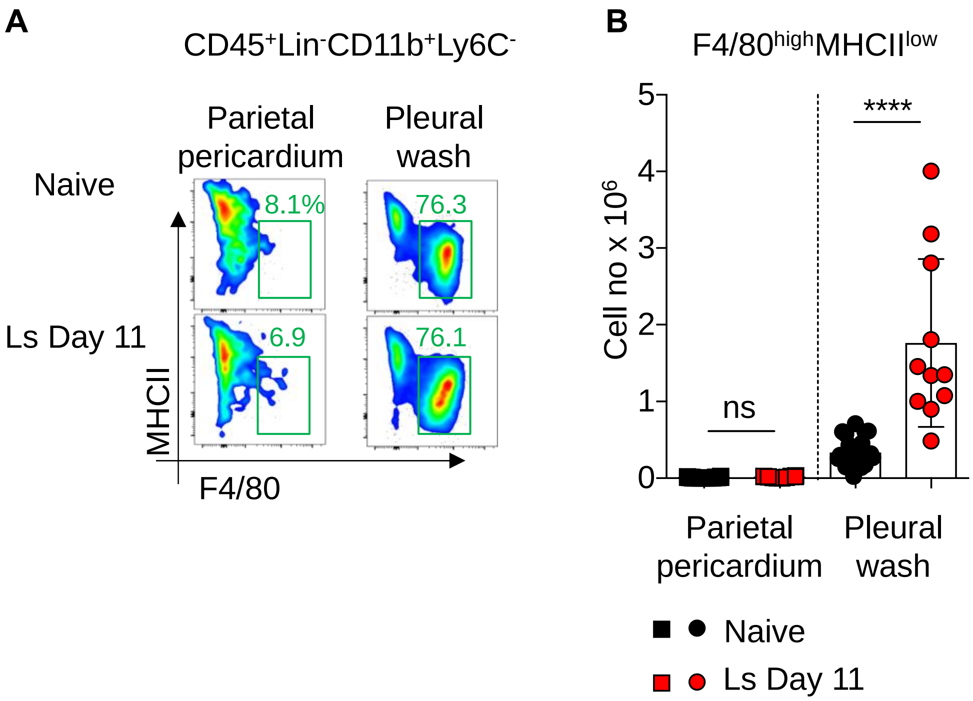


***Supplemental Figure 3: Pleural macrophages are not recruited to the parietal pericardium during pleural worm infection.*** The cellular content of the parietal pericardium and pleural wash were analyzed by flow-cytometry 11 day post-infection with Litomosoides sigmodontis (Ls). **A**: Representative flow-cytometric analysis used to identified resident pleural macrophages in the parietal pericardium (left) and pleural wash (right) of a naïve mouse (Day 0) and mice at Day 11 post-infection. Pleural macrophages were identified as Lin- CD11b+Ly6C-F4/80highMHCIIlow cells (green gate). **B**: Number of F4/80highMHCIIlow macrophages found in the parietal pericardium and pleura wash. n= 11-20 mice per group. ANOVA with Sidak's multiple comparisons test was applied. ns= non-significant, **** P=<0.0001.

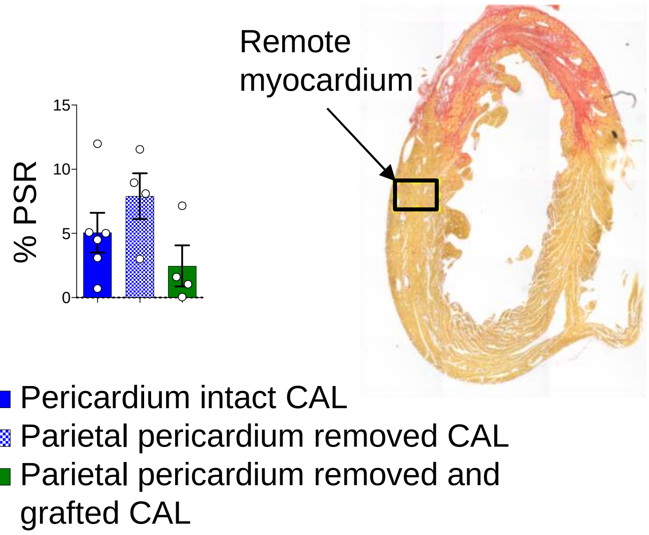


***Supplemental Figure 4: Effect of removal of the pericardium on collagen deposition in the remote myocardium.*** The pericardium was either left intact, removed or removed and re-engrafted during CAL surgeries as in Figure 4. Section of hearts were analyzed at day 7 post-MI. Representative micrographs within the left ventricular free wall showing PSR staining (red) and remote area quantified. Quantification of % positive PSR staining in remote area. Data pooled from two independent experiments n=5 mice per group. Unpaired student's T-test were performed, ns=non-significant.

| Reagent | Clone | Conjugate | Source |
|---|---|---|---|
| Armenian Hamster anti-murine TCRb | h57-597 | BV421 | Biolegend |
| Donkey anti-mouse IgM | Polyclonal | Rhodamine Red | Jackson Laboratories |
| Goat anti-rabbit | Polyclonal | biotin | Dako Cytomation |
| Rabbit anti-Ym1 | Polyclonal | | Stem Cell Technologies |
| Rat anti-I-A/I-E | M5/114.15.2 | AF780 | eBiosciences |
| Rat anti-CD11b | M1/70 | PE/Dazzle594<br>APC | Biolegend |
| Rat anti-CD19 | 6D5 | BV421 | Biolegend |
| Rat anti-CD45 | 104 | BV650<br>FITC | Biolegend |
| Rat anti-F4/80 | BM8 | PE-Cy7<br>Alexa-488 | eBiosciences |
| Rat anti-Ly6C | HK1.4 | AF700 | Biolegend |
| Rat anti-Ly6G | 1A8 | BV421 | Biolegend |
| Rat anti-Siglec-F | E50-2440 | BV421 | BD Pharmingen |

***Supplemental Table 1: List of antibodies used for flow-cytometry and immunofluorescence stainings***
